# μ-Opioid Receptor Superagonists Engage Sodium-Bound Receptor States

**DOI:** 10.64898/2025.12.01.691612

**Authors:** Alexander Powell, Nicholas W. Griggs, Jorge Iniguez-Lluhi, Henry Mosberg, John R. Traynor

## Abstract

The simple two-state conformational selection model of G-protein coupled receptor (GPCR) activation suggests that, by binding to a high affinity site, an agonist will shift receptor equilibrium in favor of active state (R*) conformations that recruit heterotrimeric G proteins over inactive state (R) conformations. Agonist binding to the μ-opioid receptor is highly sensitive to Na^+^ ions which stabilize an inactive receptor state. Higher efficacy opioid agonists, such as DAMGO and fentanyl, are sensitive to Na^+^ compared to lower efficacy ligands at the μ-opioid receptor. However, the binding of the highly potent oripavine agonists etorphine and dihydroetorphine are less sensitive to Na^+^ than expected such that the prevailing models fail to explain their pharmacology. To explain this discrepancy, experiments were performed to evaluate the binding properties and G protein activation of the highly potent agonists carfentanil, BU72, etorphine, etonitazene and similarly potent opioid peptidomimetics in comparison to the standard agonists DAMGO, fentanyl, and morphine in the presence or absence of Na^+^ or K^+^ ions. Several of the superagonists retained high affinity and potency in both ionic conditions, whereas DAMGO, fentanyl and morphine displayed enhanced binding and signaling in K^+^, compared to Na^+^ ions. These functional parameters were used to determine an intrinsic efficacy value, determined as 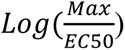. Comparison of affinity shifts with intrinsic efficacy afforded a negative correlation in which superagonists with the highest intrinsic efficacy are least sensitive to Na^+^. These data suggest that select μ-opioid receptor superagonists have high affinity for the Na^+^ bound receptor states (R) and shift these species into active receptor conformations (R*) that efficiently couple to G proteins.

**Significance Statement:** The simple theory of conformational selection suggests the binding affinities of high efficacy μ-opioid receptor ligands, such as fentanyl and DAMGO, are more sensitive to Na^+^ and guanine nucleotide which stabilize inactive receptor states than lower efficacy agonists and antagonists. Here, we show that ligands with high intrinsic efficacy (superagonists) are much less sensitive to Na^+^ and guanine nucleotide. This work demonstrates that highly potent ligands can engage a low affinity Na^+^-bound receptor state that may then convert to a receptor species that efficiently couples to G protein – i.e. a conformational induction.

## Introduction

Mu-opioid receptors (μOR) are G protein-coupled receptor (GPCR) proteins that activate heterotrimeric G proteins of the Gα_i/o_ family. The mu-opioid receptors, like other GPCRs, exist in an equilibrium of various conformational states that can be grouped into ensembles of inactive (R) and active states (R*); the R* state interacting with the G protein α-subunit. Intrinsic efficacy describes the ability of a ligand to shift the equilibrium from inactive-state conformations to active-state conformations. This parameter distinguishes ligands that prefer the “low affinity” R state (inverse agonists) from those that prefer a “high affinity” R* state (partial agonists and full agonists) and from those that display no preference (neutral antagonists). Intrinsic efficacy, in conjunction with binding affinity and pharmacokinetic properties, are the driving factors that determine the pharmacological activity of a drug for a particular receptor system.^1^

Stimulation of GTPγ^35^S binding to G proteins is a sensitive indicator of the active R* state and may be used as a predictive measure of the intrinsic efficacy of a ligand.^2^ GTPγ^35^S binding assays allow for the determination of potency (EC_50_) and relative efficacy as maximal stimulation by a ligand compared to a standard agonist. Determination of a 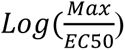 from these data is a convenient way to quantify the degree of ligand agonism.^3^ One advantage of using the GTPγ^35^S binding assay to assess ligand efficacy is that partial agonists are aptly discriminated from full agonists, since G protein activation is directly downstream of ligand-receptor binding and is less susceptible to signal amplification. On the other hand, and like other functional assays that rely on measurement of signal transduction events downstream of the receptor, results are dependent on experimental conditions.^4,5^

Agonist-mediated activation of the μOR is negatively modulated by the binding of Na^+^ ions.^6,7,8^ These ions bind deep in the receptor below the orthosteric site and form ionic interactions with transmembrane domains 2, 3, and 7, holding the receptor in an inactive set of conformations.^9^ Thus, agonists and Na^+^ ions work in opposition; agonists shift the equilibrium toward active R* ensembles whereas Na^+^ ions shift the equilibrium toward inactive R ensembles.^9,10,11^ This negative cooperativity has been long known at μOR, in which the addition of Na^+^ ions with guanine nucleotide to uncouple G proteins, decreases the binding affinity (*K_i_*) of opioid agonists.^6,12^ Higher efficacy opioid agonists show a greater decrease in binding affinity upon the addition of Na^+^ and guanine nucleotide compared to lower efficacy agonists and antagonists display no shift in affinity.^6,13^ There appears to be a positive association between the decrease in the *K_i_* of an agonist by Na^+^ (i.e. sodium sensitivity) and the intrinsic efficacy of an opioid. However, two very potent opioid agonists, etorphine and dihydroetorphine, demonstrate significantly reduced sodium sensitivity which questions the validity of the observed positive association between sodium sensitivity and intrinsic efficacy at μOR.^13^

The effects of sodium on μOR activation cannot be treated as uniform or universal. Sodium may stabilize multiple inactive conformations, but its allosteric influence is probe-dependent and varies with the specific agonist engaged, based largely on agonist efficacy. Different ligands couple to the receptor in distinct ways that either disrupt or accommodate sodium binding.^13,14^ For superagonists, it is possible that receptor engagement does not require displacement of the Na^+^ ion; instead, these ligands may interact with Na^+^-bound states leading to displacement of Na^+^ and receptor activation, i.e. conformational induction or induced fit. Alternatively, superagonists bind and stabilize Na^+^-bound receptor ensembles that are still competent to recruit G protein. Thus, receptor activation may proceed through several parallel trajectories, with some involving sodium displacement and others proceeding through Na⁺-bound “active-like” states that still facilitate G protein coupling. These observations suggest that the role of Na^+^ in μOR activation should not be considered strictly inhibitory but instead may include Na⁺-bound states that remain competent to interact with high-efficacy agonists.

Here, we report the determination of intrinsic efficacy for a series of potent opioids including carfentanil, BU72, etorphine and etonitazene, together with a series of highly potent opioid peptidomimetics, and compare them with standard opioids morphine, DAMGO and fentanyl (**Figure 1**). By evaluating the affinity of these ligands in the absence and presence of Na^+^ and guanine nucleotide and calculating the difference in affinity between the two conditions (i.e. “affinity shift”), we show that for highly potent opioid agonists, reduced Na^+^ sensitivity correlates significantly with higher intrinsic efficacy. The model of conformational selection would predict the compounds to be neutral antagonists or low efficacy agonists, as they demonstrate small affinity shifts and recognize different affinity states (inactive and active) in a near equal manner. Instead, we suggest a conformational induction model in which highly potent agonists bind to a low-affinity state of μOR in the presence of Na^+^ ions and shift the receptor into a high-affinity state capable of recruiting G protein.

**Figure 1:**
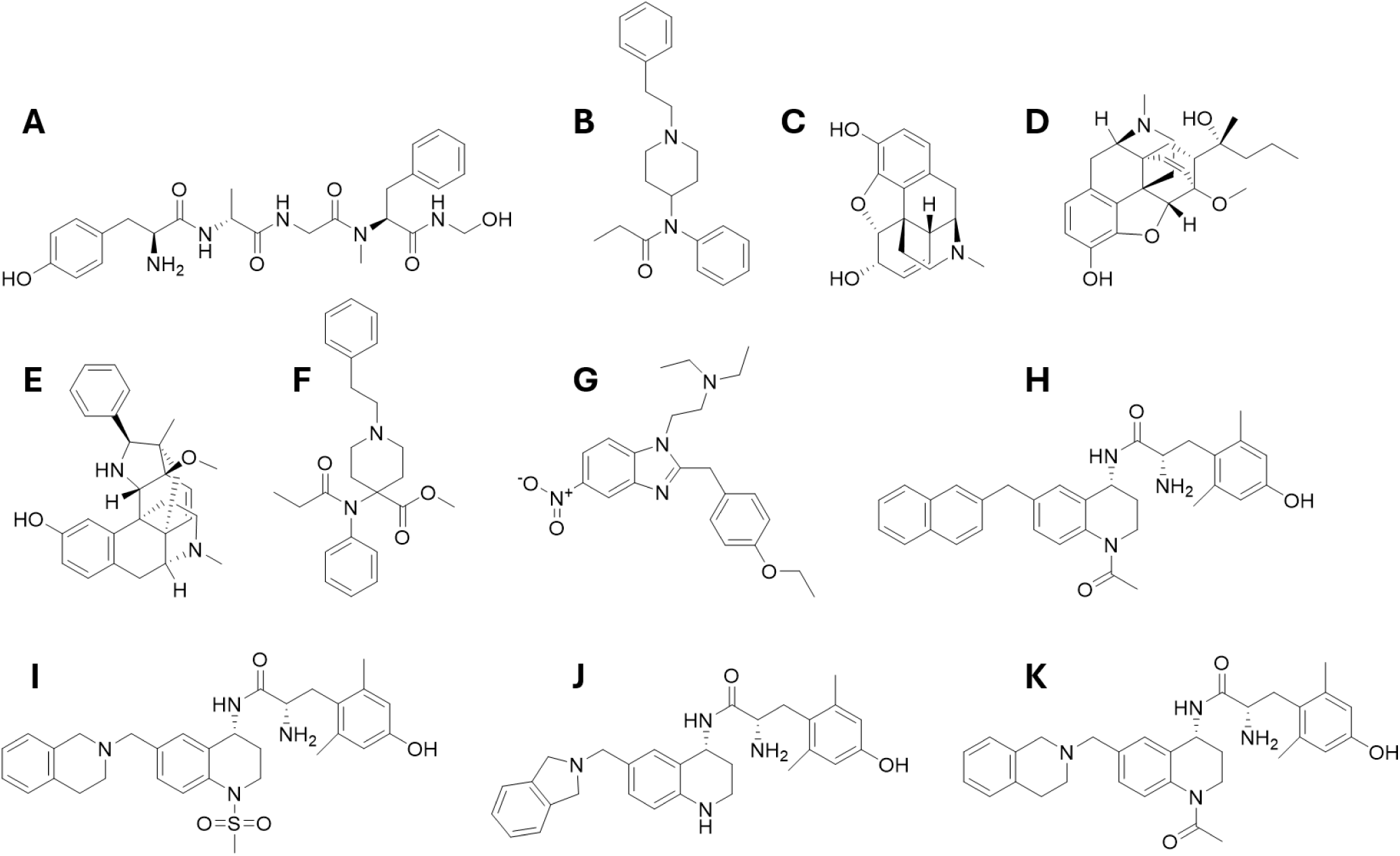
Chemical Structures of Opioid Agonists and Peptidomimetics. (A) DAMGO, (B) Fentanyl, (C) Morphine, (D) Etorphine, (E) BU72, (F) Carfentanil, (G) Etonitazene, (H) AAH8, (I) AFN42, (J) AMB46, and (K) AMB47

## Methods

### Materials and Reagents

We have previously reported the synthesis and pharmacological evaluation of novel opioid peptidomimetic compounds.^15,16,17,18,19,20,21^ AAH8 (compound 14a) was synthesized by Aubrie A. Harland^16^, AFN42 (compound 4d) was synthesized by Anthony F. Nastase^21^, AMB46 (compound 10n) and AMB47(Compound 10m) were synthesized by Aaron M. Bender.^15^ All chemicals, unless otherwise specified, were purchased from Sigma Aldrich. Radiolabeled [^3^H]-diprenorphine (DPN) and radiolabeled guanosine-5′-O-(3-[^35^S] thio)-triphosphate (GTPγ^35^S) were purchased from PerkinElmer Life Sciences. All tissue culture media, penicillin-streptomycin, geneticin (G148), trypsin, and fetal bovine serum were purchased from Gibco LifeSciences.

### Cell Lines

The tissue culture and maintenance of C6 rat glioma cells stably transfected with rat μOR was performed as previously described.^21^ C6 cells were grown to confluence at 37 °C in 5 % CO_2_ in Dulbecco’s modified Eagle media (DMEM) containing 10 % fetal bovine serum (FBS), 5 % penicillin/streptomycin (P/S), and 400 ug/mL G418. CHO cells stably expressing wild-type human μOR were grown to confluence at 37 °C in 5 % CO_2_ in Dulbecco’s modified Eagle media (DMEM) containing 10 % fetal bovine serum (FBS), 1 % penicillin/streptomycin (P/S), and 400 ug/mL G418.

### Cell Membrane Preparation

Cell membrane homogenates from C6 and CHO cells were prepared by washing confluent cells three times with phosphate buffered saline (pH 7.4), after which they were then detached using harvesting buffer (20 mM HEPES, 150 mM NaCl, 0.68 mM EDTA, pH7.4) and pelleted by centrifugation at 200 x g for 3 min at room temperature. The pellet was resuspended in ice-cold 50 mM Tris buffer (pH 7.4) and homogenized using a Tissue Tearor (Biospec Products, Inc.). The cell lysate was centrifuged at 20,000 x g at 4 °C for 20 minutes, after which the pellet was then resuspended on ice cold 50 mM Tris (pH7.4), homogenized, and centrifuged once more at 20000 x g at 4 C for 20 minutes. The final pellet was resuspended in 50 mM Tris (pH 7.4) using a glass dounce homogenizer and then dispensed into aliquots and stored at -80 °C. The final protein concentration was determined by performing a BCA protein assay (Thermo Scientific Pierce) using bovine serum albumin as the standard.

### Competition Radioligand Binding Assays

^3^H-diprenorphine competition binding assays were performed in 500μL reactions in 96-well plates using the cell membrane homogenates described above. Briefly, cell membranes [5-20 μg] were incubated in 50 mM Tris buffer containing 0.2 nM ^3^H-Diprenorphine, and various concentrations of opioid compound for 90 minutes in a shaking water bath at room temperature to reach equilibrium. Nonspecific binding was determined using 10 µM of the opioid antagonist, naloxone. The reaction was terminated by vacuum filtration onto glass microfiber GF/C filters using a tissue harvester (Brandel, Gaithersburg, MD) and washed with 50 mM Tris buffer. Filters were dried and following the addition of EcoLume scintillation cocktail, bound radioactivity was quantified in a MicroBeta 2450 Liquid Scintillation and Luminescence Counter (PerkinElmer).

Radioligand binding assays in Na^+^ or K^+^ containing buffers were performed in 200 μL reactions in 96-well plates using the cell membrane homogenates described above, Briefly, cell membranes [5-20 μg] were incubated in Na^+^ or K^+^ containing buffers (50 mM Tris, 100 mM NaCl or 100 mM KCl, 5 mM MgCl2, 1 mM EDTA, pH 7.4) or (50 mM Tris, 300 mM NaCl, 5 mM MgCl2, 1 mM EDTA, pH 7.4) containing 10 μM cold GTPγS, and various concentrations of opioid compound for 90 minutes in a shaking water bath at room temperature. Nonspecific binding was determined using 10 µM naloxone. The reaction was terminated by vacuum filtration onto glass microfiber GF/C filters using a tissue harvester and washed with Na^+^ containing buffer, and the filters were processed as described above.

### Stimulation of GTPγ^35^S Binding Assays

Stimulation of GTPγ^35^S binding experiments were performed in 200 μL reactions in 96-well plates using cell membrane homogenates prepared as described above. Briefly, cell membranes [10-20 μg] were incubated in either Na^+^ or K^+^ containing buffer [50 mM Tris, 100 mM NaCl or 100mM KCl, 5 mM MgCl2, 1 mM EDTA, pH 7.4] containing 0.1 nM guanosine-5′-O-(3-[^35^S]thio)triphosphate (GTPγ^35^S) and 30 μM guanosine 5’-diphosphate (GDP) and various concentrations of opioid compound or peptidomimetic for 1 hour in a shaking water bath at 25 °C. Basal stimulation of GTPγ^35^S binding was determined by incubation in the absence of any ligand and maximal stimulation of was defined by using 10 μM of the standard mu opioid receptor agonist, DAMGO ([D-Ala^2^,N-MePhe^4^,Gly-ol]-enkephalin). The reaction was terminated by vacuum filtration onto glass microfiber GF/C filters using a tissue harvester and washed with Na^+^ containing buffer, and filters were processed as described above.

## Results

### GTPγ^35^S Binding

As evaluated by concentration response curves of GTPγ^35^S binding in C6 μOR membranes, etorphine, BU72, carfentanil and etonitazene were seen to be very potent agonists with EC_50_ values in the low nanomolar and sub-nanomolar range **(Table 1 and Figure 2)**. In addition, we have previously described a series of highly potent peptidomimetics that bind to μOR with exceptional affinity.^15,16,21^ These compounds (H-K in **Figure 1**) are at least 70 times more potent than the standard opioid peptide DAMGO, fentanyl, or morphine with potencies that are similar to carfentanil and etonitazene. All compounds demonstrated high relative efficacy, similar to DAMGO, as evaluated by maximal stimulation of GTPγ^35^S binding. The fractional maximal response (i.e. “Max”) compared to DAMGO (Max = 1) was equivalent for all ligands.

**Figure 2:**
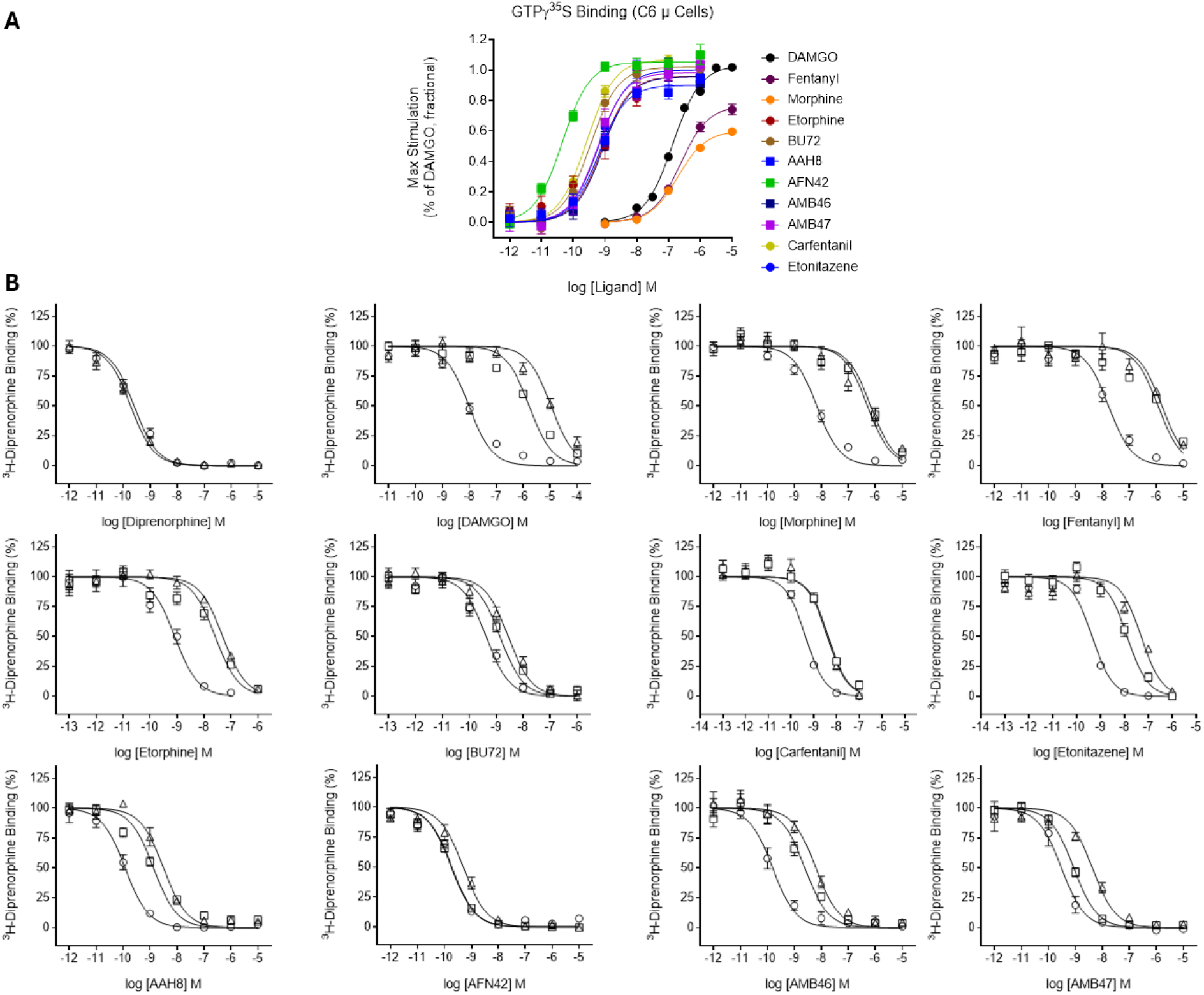
Stimulation of GTPγ^35^S Binding and Inhibition of ^3^H-Diprenorphine Binding Concentration-Response Curves in C6 μ Cells. (A) GTPγ^35^S binding assays were performed using membranes from C6 cells with various concentrations of opioid agonists. Maximal stimulation of GTPγ^35^S binding was produced by 10 μM of the reference agonist, DAMGO. (B) Radioligand competition binding assays were performed in the following buffers: 50 mM Tris (circles), 50 mM Tris, 100 mM NaCl, 5 mM MgCl_2_ 1 mM EDTA, 10 μM GTPγS (squares), or 50 mM Tris, 300 mM NaCl, 5 mM MgCl_2_ 1 mM EDTA, 10 μM GTPγS (triangles). These assays used 20 μg membrane protein from C6 cells, 0.2 – 0.5 nM ^3^H-Diprenorphine, and various concentrations of the following opioid ligands: Diprenorphine, DAMGO, Fentanyl, Etorphine, BU72, Carfentanil, Etonitazene, AAH8, AFN42, AMB46, AMB47. All experiments were performed in duplicate or quadruplicate (n ≥ 3).

**Table 1:** Stimulation of GTPγ^35^S Binding and Inhibition of ^3^H-Diprenorphine Binding by Opioid Ligands in C6 μ Cells. Stimulation of GTPγ^35^S binding at the μOR was performed using membranes from C6 cells with various concentrations of opioid agonists. Maximal stimulation of GTPγ^35^S binding was determined by 10 μM of the reference agonist, DAMGO. Effective concentrations (EC_50_) and the fractional maximal response (Max Stimulation) were determined by nonlinear regression analysis of concentration-response curves from GTPγ^35^S binding assays. Agonism (Log [Max/EC50]) was calculated using EC_50_ and max stimulation values for each opioid agonist. ^3^H-Diprenorphine binding assays were performed using 0.2 – 0.5 nM of the radioligand in the following three buffers: **1.** 50 mM Tris buffer, **2.** 50 mM Tris, 100 mM NaCl, 5 mM MgCl_2_ 1 mM EDTA, and 10 μM GTPγS, or **3.** 50 mM Tris, 300 mM NaCl, 5 mM MgCl_2_ 1 mM EDTA, and 10 μM GTPγS. The affinity shift was determined for 100 mM Na^+^ and 300 mM Na^+^ conditions. Data are represented as mean plus 95% confidence intervals and all experiments were performed in duplicate or quadruplicate (n ≥ 3).

| C6 $\mu$ Cells | | GTPγ<sup>35</sup>S Binding | | <sup>3</sup>H-Diprenorphine Binding | | | | |
| --- | --- | --- | --- | --- | --- | --- | --- | --- |
| Compound | EC<sub>50</sub> (nM) | Max Stimulation<br>(% of DAMGO, fractional) | Agonism<br>Log $\frac{\text{Max}}{\text{EC}_{50} \text{ (M)}}$ | K<sub>i</sub> (nM)<br>50 mM Tris | K<sub>i</sub> (nM)<br>+ 100mM Na<sup>+</sup><br>+10μM GTPγS | Affinity Shift<br>K<sub>i</sub> 100mM Na +<br>K<sub>i</sub> 50mM Tris | K<sub>i</sub> (nM)<br>+ 300mM Na<sup>+</sup><br>+10μM GTPγS | Affinity Shift<br>K<sub>i</sub> 300mM Na +<br>K<sub>i</sub> 50mM Tris |
| DAMGO | 42.2<br>(39.0-47.9) | 1 | 7.38 | 4.71<br>(3.63-6.13) | 618<br>(421-929) | 131 | 5840<br>(4040-8470) | 1240 |
| Fentanyl | 233<br>(178-308) | 0.76<br>(0.72-0.80) | 6.51 | 8.33<br>(5.74-12.3) | 433<br>(239-780) | 52.0 | 873<br>(488-1590) | 105 |
| Morphine | 195<br>(148-260) | 0.60<br>(0.57-0.63) | 6.49 | 3.32<br>(2.47-4.42) | 254<br>(169-377) | 76.5 | 280<br>(155-485) | 84.3 |
| Diprenorphine | - | - | - | 0.12<br>(0.094-0.16) | - | - | 0.10<br>(0.077-0.13) | 0.83 |
| Etorphine | 0.79<br>(0.42-1.39) | 0.96<br>(0.89-1.04) | 9.08 | 0.40<br>(0.24-0.66) | 9.66<br>(6.02-15.4) | 24.2 | 26.6<br>(19.7-35.6) | 66.5 |
| BU72 | 0.35<br>(0.24-0.51) | 1.02<br>(0.96-1.08) | 9.46 | 0.20<br>(0.14-0.29) | 0.50<br>(0.33-0.77) | 2.5 | 1.48<br>(0.96-2.33) | 7.4 |
| Carfentanil | 0.37<br>(0.28-0.49) | 1.06<br>(1.01-1.11) | 9.46 | 0.17<br>(0.13-0.23) | 2.05<br>(1.37-3.03) | 12.1 | 2.36<br>(1.63-3.40) | 13.9 |
| Etonitazene | 0.63<br>(0.38-1.03) | 1.0<br>(0.92-1.08) | 9.20 | 0.18<br>(0.14-0.22) | 5.94<br>(4.19-8.46) | 33 | 29.8<br>(19.3-45.1) | 166 |
| AAH8 | 0.64<br>(0.43-0.95) | 0.9<br>(0.85-0.95) | 9.15 | 0.058<br>(0.043-0.079) | 0.50<br>(0.37-0.68) | 8.6 | 1.70<br>(1.25-2.31) | 29.3 |
| AFN42 | 0.058<br>(0.043-0.081) | 1.06<br>(1.01-1.11) | 10.26 | 0.093<br>(0.072-0.12) | 0.067<br>(0.054-0.083) | 0.72 | 0.27<br>(0.19-0.36) | 2.9 |
| AMB46 | 0.86<br>(0.53-1.37) | 0.96<br>(0.89-1.03) | 9.04 | 0.077<br>(0.052-0.11) | 0.91<br>(0.60-1.42) | 11.8 | 3.39<br>(2.17-5.23) | 44 |
| AMB47 | 0.59<br>(0.37-0.92) | 0.98<br>(0.91-1.05) | 9.22 | 0.15<br>(0.097-0.28) | 0.32<br>(0.27-0.38) | 2.1 | 2.29<br>(1.65-3.15) | 15.3 |

### Agonist Binding Affinities

It has been known for decades that physiologically relevant Na^+^ ion concentrations decrease the binding affinity of agonists to μOR.^22^ Radioligand competition binding assays were performed using membranes from C6 cells expressing the rat μOR in the presence or absence of Na^+^ ions and guanine nucleotide (GTP**γ**S) to induce R and R* receptor states, respectively. Difference in binding affinities between the two conditions were given as affinity shifts (**Table 1 and Figure 2**). In 50 mM Tris, the superagonists carfentanil, BU72, etorphine and etonitazene bound with approximately 10-fold higher affinity than the classical agonists DAMGO, morphine and fentanyl. The peptidomimetics exhibited similar binding affinities, with AAH8 displaying the highest affinity.

Binding affinities of DAMGO, fentanyl, and morphine to μOR decreased in the presence of 100 mM Na^+^ and GTP**γ**S, producing affinity shifts of 131, 52 and 77-fold, respectively (**Table 1**). Of the superagonists, etorphine and etonitazene displayed the greatest affinity shifts of 24 and 33-fold, respectively. Carfentanil, AAH8 and AMB47 all displayed 10-fold decreases in affinity, whereas BU72, AFN42 and AMB47 displayed much smaller shifts or no shift.

Since the superagonists displayed a less than expected affinity shift in the presence of 100 mM Na^+^ and GTP**γ**S, we utilized a supraphysiological concentration of Na^+^ to determine whether the affinity shift for each superagonist was saturated (**Table 1**). In the presence of 300 mM Na^+^ ions and GTP**γ**S, an affinity shift of 1,240-fold was observed for DAMGO, whereas fentanyl and morphine displayed smaller shifts of 105 and 84-fold, respectively. Of the highly potent agonists, only etonitazene produced an affinity shift greater than 100-fold. Etorphine, AAH8 and AMB46 had comparable affinity shifts ranging from 29 to 67-fold. Carfentanil and AMB47 displayed smaller shifts of 14 and 15-fold, respectively. Two compounds exhibited shifts of less than 10-fold, these ligands being BU72 (7-fold) and AFN42 (3-fold).

### Comparison of the Effect of Na^+^ and K^+^ Ions on Agonist-Mediated GTPγ^35^S Binding and Ligand Affinity

Since we compared agonist binding in Na^+^ and GTP**γ**S to binding in 50 mM Tris-HCl buffer alone, we needed to evaluate the role of ionic strength on the binding and functional measures. To do this, we replaced NaCl with the same concentration of KCl and examined two superagonists, carfentanil and AFN42. The conserved class A GPCR allosteric sodium site is defined by a tight coordination network involving Asp^2.50^ and multiple other polar residues. This geometry selectively accommodates Na^+^ and is not observed with larger monovalent cations, like K^+^, in solved crystal structures.^23^ Therefore, the larger K^+^ ion would not be expected to efficiently hold the receptor in an inactive conformation. Consequently, in the GTP**γ**^35^S assay the standard opioid agonist DAMGO exhibited 4-fold greater potency in 100 mM KCl relative to 100 mM NaCl (11 nM (8.4 – 16 nM) vs. 42 nM (39 – 48 nM) respectively), consistent with previous evidence that Na^+^ stabilizes an inactive receptor (R) conformations and so inhibits G protein coupling (**Figure 3**).^6,7,9^ In support, diprenorphine exhibited partial agonist activity in the 100 mM KCl condition. However, the superagonist tested AFN42 demonstrated increased potency in the presence of Na^+^ compared to K^+^ (AFN42 in NaCl: 0.058 nM (0.043 – 0.081 nM) vs. in KCl: 0.25 nM (0.19 – 0.35 nM) and carfentanil near equivalent potency (carfentanil in NaCl: 0.37 nM (0.28 – 0.49 nM) vs in KCl (0.19 nM (0.15 – 0.25 nM). This suggests that Na^+^ does not universally prevent agonist activity unlike previously thought to be true.

**Figure 3:**
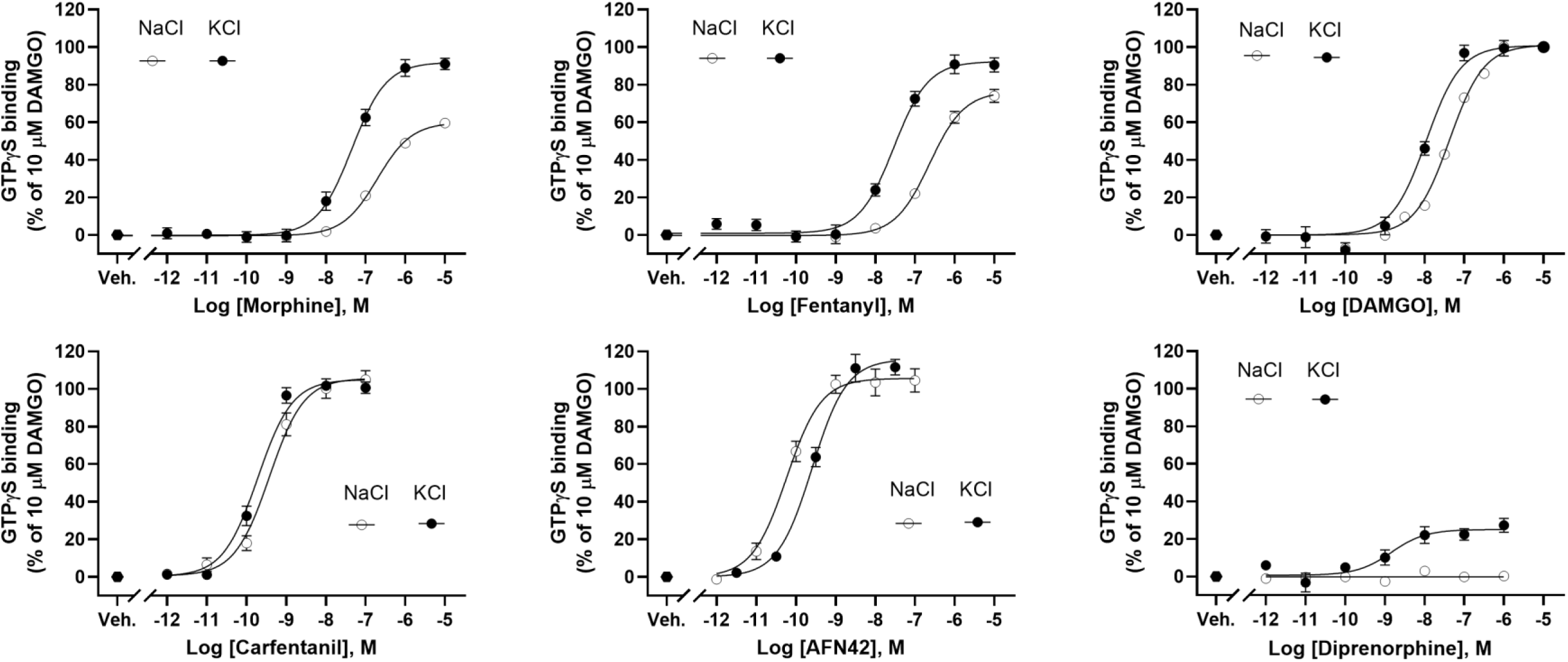
Stimulation of GTPγ^35^S Binding in 100 mM NaCl and KCl buffer. Stimulation of GTPγ^35^S binding at the μOR was performed using membranes from C6 cells with increasing concentrations of opioid agonists. Maximal stimulation of GTPγ^35^S binding was determined by 10 μM of the reference agonist, DAMGO. Effective concentrations (EC_50_) were determined by nonlinear regression analysis of concentration-response curves from GTPγ^35^S binding assays. Data are represented as mean plus 95% confidence intervals and all experiments were performed in duplicate (n ≥ 3).

To evaluate the contribution of Na^+^ ions and GTP**γ**S on ligand-receptor interactions, we compared the above results with radioligand binding assays in the presence of 100 mM KCl and GTP**γ**S. These latter conditions lead to a smaller rightward shift in affinity, consistent with the inability of K^+^ ions to replace Na^+^ ions and stabilize a low affinity state (**Table 2**). In contrast, AFN42 displayed a greater rightward shift in 100 mM KCl (18.1-fold) than it did in the presence of 100 mM NaCl (0.7-fold) or even 300 mM NaCl (2.9-fold) (**Table 2**). Similarly, carfentanil displayed no significant change in affinity when examined in 100 mM KCl (K_i_: 1.2 nM), 100 mM NaCl (K_i_: 2.1 nM), or 300 mM NaCl (K_i_: 2.4 nM).

**Table 2.** Inhibition of ^3^H-Diprenorphine Binding in 100 mM KCl. Radioligand competition binding assays were performed in 50 mM Tris, 100 mM KCl, 5 mM MgCl_2_ 1 mM EDTA, 10 μM GTPγS. These assays used 20 μg membrane protein from C6 cells, 0.2 – 0.5 nM ^3^H-Diprenorphine, and various concentrations of opioid ligand. Experiments were performed in duplicate or quadruplicate (n ≥ 3) and are presented as mean with 95% confidence intervals.

| C6 $\mu$ Cells | <sup>3</sup>H-Diprenorphine Binding | | |
| --- | --- | --- | --- |
| Compound | K<sub>i</sub> (nM)<br>50 mM Tris | K<sub>i</sub> (nM)<br>+ 100mM K<sup>+</sup><br>+10μM GTPγS | Affinity Shift<br>$\frac{\text{Ki 100mM K} +}{\text{Ki 50mM Tris}}$ |
| DAMGO | 4.71<br>(3.63-6.13) | 111<br>(66.8-187) | 23.6 |
| Fentanyl | 8.33<br>(5.74-12.3) | 218<br>(140-344) | 26.2 |
| Morphine | 3.32<br>(2.47-4.42) | 150<br>(93.9-246) | 45.2 |
| Diprenorphine | 0.12<br>(0.094-0.16) | 1.96<br>(1.29-2.94) | 16.3 |
| Carfentanil | 0.17<br>(0.13-0.23) | 1.19<br>(0.81-1.76) | 7 |
| AFN42 | 0.093<br>(0.072-0.12) | 1.68<br>(1.02-2.86) | 18.1 |

These data suggest that Na^+^ is not always inhibitory to agonist binding and indicates superagonists are capable of binding to a Na^+^-bound receptor state. It may then convert this receptor species into a Na^+^-free state that can then efficiently couple to G proteins.

### Agonist Binding in CHO Cells Expressing Human μOR

Since the above data was obtained in C6 glioma cells expressing the rat μOR, we performed experiments in CHO cells expressing human μOR (CHO-μOR). Radioligand binding experiments were performed in CHO-μOR cell membranes in the absence or presence of 100 mM Na^+^ ions and GTP**γ**S to confirm that the affinity shifts observed were not specific to the rat μOR or dependent on the cell type/receptor environment (**Table 3**). Three ligands were examined – DAMGO, BU72 and carfentanil. In the presence of 100 mM Na^+^ and GTP**γ**S, DAMGO showed a substantial decrease in binding affinity, resulting in a shift of 810-fold, larger than the 131-fold shift observed in the C6 μOR cells. Both BU72 and carfentanil exhibited smaller shifts of 3 and 35-fold, respectively, although these shifts were greater than observed in the C6 cells. The differences in affinity shifts observed between ligands and cell types could be due to differences between the human and rat μOR, as there are subtle variations in amino acid sequences which have been shown can lead to differences in ligand binding affinities.^24^ Additionally, their cellular environments (hamster ovary versus rat glioma cell) and access to different accessory proteins on the membranes may also be of influence. Nevertheless, the same pattern is observed across the two species of μOR and cell types.

**Table 3:**
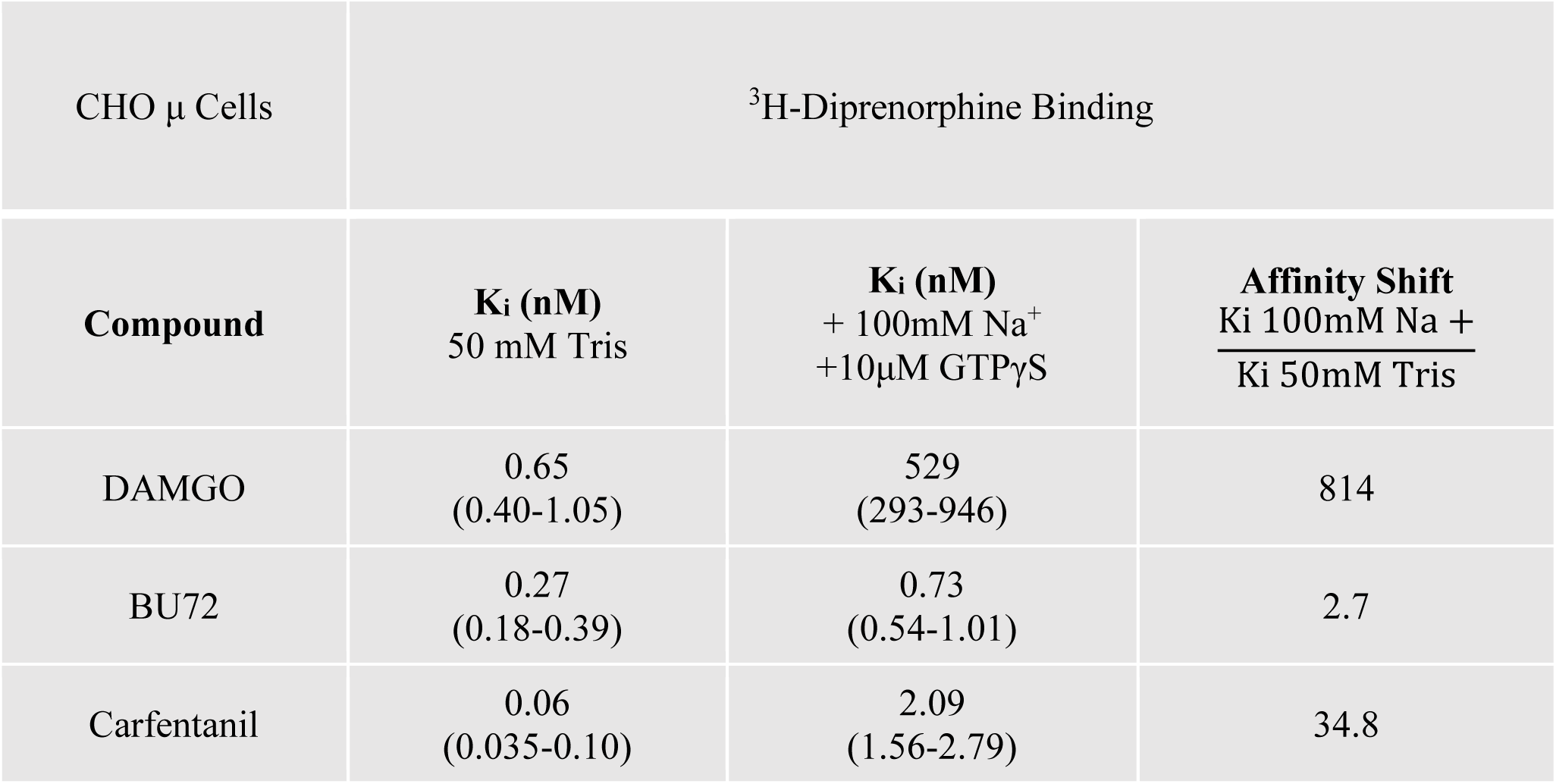
Inhibition of ^3^H-Diprenorphine Binding by Opioid Ligands in CHO μ Cells. ^3^H-Diprenorphine binding assays were performed using 0.2 – 0.5 nM of the radioligand in the following two buffers: **1.** 50 mM Tris buffer, and **2.** 50 mM Tris, 100 mM NaCl, 5 mM MgCl_2_ 1 mM EDTA, and 10 μM cold GTPγS. The affinity shift was determined for 100 mM Na^+^ conditions. Data are represented as mean plus 95% confidence intervals and all experiments were performed in duplicate (n ≥ 3).

| CHO $\mu$ Cells | $^3\text{H}$-Diprenorphine Binding | | |
| --- | --- | --- | --- |
| Compound | **K<sub>i</sub> (nM)**<br>50 mM Tris | **K<sub>i</sub> (nM)**<br>+ 100mM Na<sup>+</sup><br>+10μM GTPγS | **Affinity Shift**<br>$\frac{\text{K}_i \text{ 100mM Na} +}{\text{K}_i \text{ 50mM Tris}}$ |
| DAMGO | 0.65<br>(0.40-1.05) | 529<br>(293-946) | 814 |
| BU72 | 0.27<br>(0.18-0.39) | 0.73<br>(0.54-1.01) | 2.7 |
| Carfentanil | 0.06<br>(0.035-0.10) | 2.09<br>(1.56-2.79) | 34.8 |

### Correlating Intrinsic Efficacy with Affinity Shifts in NaCl Conditions

The level of agonism was determined for each opioid agonist by calculating the 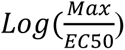 from GTPγ^35^S binding assays performed in 100 mM NaCl (as described in Kenakin, 2017).^3^ The value for AFN42 (10.26) is much greater than that of DAMGO (7.38). Since these parameters are on a logarithmic scale, the difference in agonism between AFN42 and DAMGO (10.26 – 7.38 = 2.88) is a 760-fold difference. Since the superagonists tested give the same maximal response the difference in agonism is attributable to the difference in EC_50_. The agonism value for each compound was then plotted in comparison to the affinity shift seen in the presence of Na^+^ ions and GTP**γ**S (**Figure 4**). This plot shows that for full agonists in the GTP**γ**^35^S assay, agonism and affinity shift are negatively correlated in both the 100 mM Na^+^/GTP**γ**S conditions (r^2^ = 0.63, p-value < 0.003) and the 300 mM Na^+^/GTP**γ**S conditions (r^2^ = 0.45, p-value = 0.02). Thus, both the level of agonism and Na⁺-dependent affinity shifts serve as complementary and predictive readouts of superagonism at the receptor.

**Figure 4:**
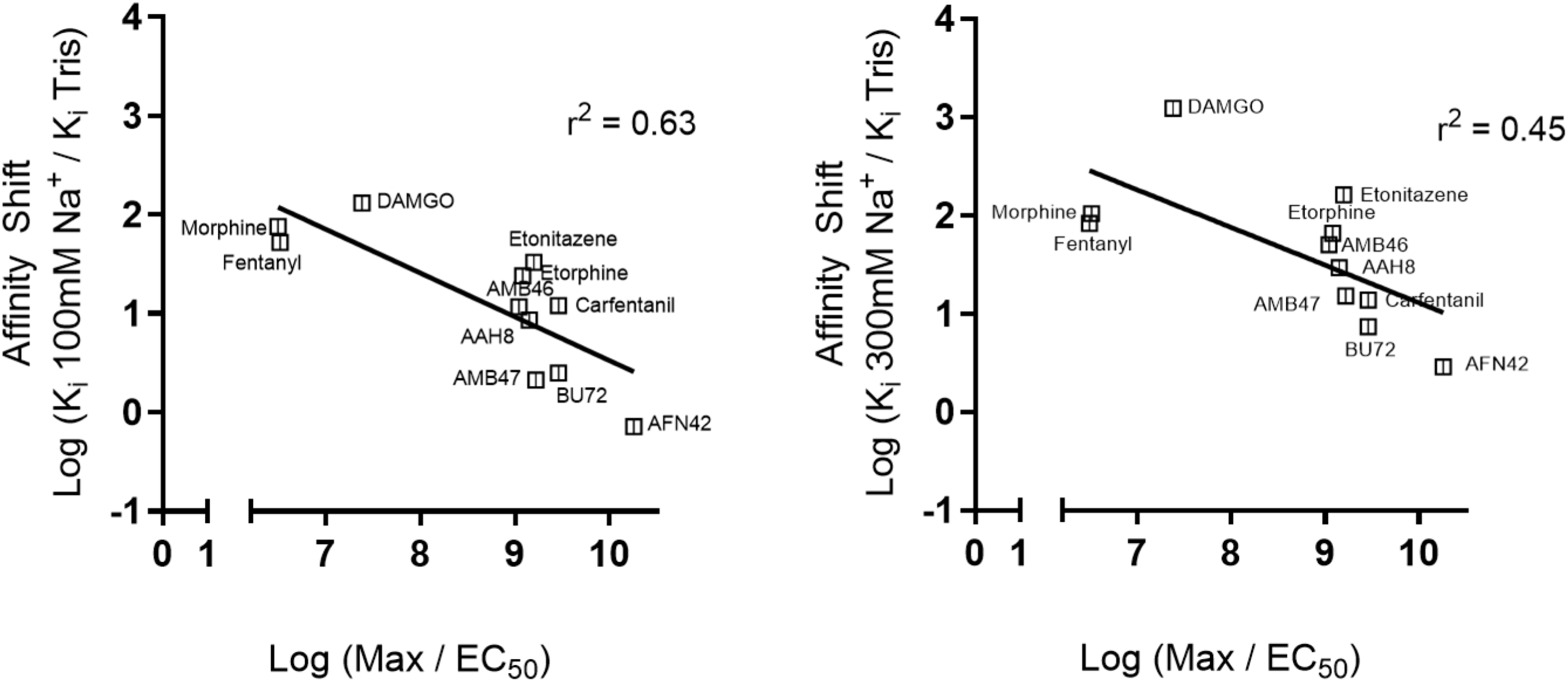
Intrinsic Efficacy Plots for C6 μ. Agonism (Log [Max/EC_50_]) was determined from stimulation of GTPγ^35^S binding assays and Affinity Shift (Log [Ki, 100 mM Na^+^ / K_i_ 50 mM Tris] in **(A)** or Log [Ki, 300 mM Na^+^ / K_i_ 50 mM Tris] in **(B)** was determined from ^3^H-Diprenorphine competition binding assays. Data was plotted for each of the following ligands: DAMGO, Fentanyl, Morphine, Etorphine, BU72, Carfentanil, Etonitazene, AFN42, AAH8, AMB46, and AMB47. r^2^ = 0.63, p** = 0.003 **(A)**; r^2^ = 0.45, p* = 0.02 **(B)**

## Discussion

In this study, we investigate the potency and binding affinity of a series of highly potent agonists (superagonists) at μOR using GTPγ^35^S assays, and radioligand competition binding assays in the absence or presence of Na^+^ ions and guanine nucleotide (GTPγS). Na^+^ ions act as negative allosteric modulators of the receptor, stabilizing the inactive R resting state. Similarly, guanine nucleotides prevent the G protein from stabilizing the active R* state by binding to the Gα subunit of the G protein heterotrimer. Agonists preferentially bind to the active R* receptor state, exhibiting higher affinity for R* than for R. Under the conformational selection model, this binding shifts the receptor population or equilibrium toward the active R* state. However, our data comparing differences in agonist affinities in two ionic conditions with the GTPγ^35^S functional data revealed that superagonists may be far less sensitive to Na^+^ ions and guanine nucleotide than predicted. Although the conformational selection model predicts the action of most types of ligands (such as endogenous agonists and agonists with similar kinetic properties, inverse agonists, partial agonists and antagonists), our data suggests it falls short in explaining the action of extremely potent opioid agonists, which appear able to function through a conformation induced or induced-fit model, binding to an inactive Na^+^-bound R states and converting these to active R* states.

Binding assays performed in conditions in which Na^+^ ions were replaced with K^+^ ions to maintain osmolarity highlighted the difference in behavior seen between superagonists and less potent agonists. DAMGO and morphine bind more tightly to μOR in the presence of 100 mM KCl and GTPγS, consistent with the absence of Na^+^-mediated allosteric inhibition. However, AFN42 exhibits a paradoxical increase in binding affinity in the presence of 100 mM or even 300 mM Na^+^ ions and GTPγS compared to its affinity in 100 mM K^+^ ions with GTPγS. This suggests that Na^+^ occupancy of its allosteric site on μOR may even favor the binding of some superagonists, supporting a model in which these agonists access a Na^+^-bound receptor conformation that is far less accessible to conventional, less potent agonists. Thus, Na^+^ ions do not exhibit a uniform inhibitory effect across all agonists, rather the role of Na^+^ as a negative allosteric modulator can be in part dependent on agonist affinity. In support of this, at the delta-opioid receptor (δOR), binding of the highly potent agonist BW373U86 has been reported as insensitive to Na^+^ ions and guanine nucleotide, compared to a peptide agonist.^25^ However, high-affinity superagonists are not universally agnostic to the effects of Na^+^ ions as seen in the aforementioned study with the peptide agonist DSLET and in our own study with etonitazene. It may reflect a specific interaction, or a coordinated interaction network, that mediates this Na^+^ insensitivity.

A recent structural and computational study of μOR partial, full and superagonists demonstrate that G protein activation proceeds through a series of intermediate conformational states that are separated by energetic barriers, and that the differences in agonist efficacy arise from the kinetics of traversing these steps rather than from distinct activation pathways.^26^ Superagonists increase receptor dynamics, allowing for a substantially more rapid transition towards active ternary complexes relative to full and partial agonists, where partial agonists stabilize more rigid intermediate conformations that struggle to overcome these energy barriers needed to initiate fully active complexes. Consistent with this, molecular dynamics simulations initiated from the inactive μOR demonstrate that the superagonist lofentanil can drive spontaneous transitions into an active receptor-G protein complex in a subset of trajectories, further indicating that superagonists can bind to inactive receptors and markedly reduce the energetic barrier required for activation.^26^

The Na^+^ ion binding site on class A GPCRs comprising of Asp^2.50^, Asn^1.50^, Ser^3.39^, Asn^7.45^, Ser^7.46^, Asn^7.49^ and Tyr^7.53^ is highly conserved across GPCRs.^9^ For example, Asp^2.50^, is found in 90% of non-olfactory class A GPCRs, including the opioid receptor family.^9,27,28^ Mutating Asp^2.50^ causes different functional outcomes depending on the GPCR. For example, with the rat μOR, a D114(2.50) N mutation reduces binding affinities, potencies and efficacies of both DAMGO and morphine, as well as the binding affinity of the peptide antagonist, CTAP.^29^ Mutagenesis studies of this residue in the closely related δOR transform antagonists into potent beta-arrestin-biased agonists, suggesting an important role in mediating the R-R* equilibria.^31^ Similar experiments have been performed in a range of Class A GPCRs and lead to a variety of different pharmacological effects, including reduced downstream signaling and receptor internalization, increases in agonist affinity with no change in antagonist affinity, no change in agonist affinity, and changes in antagonist affinity while abolishing G protein signaling.^31–34^ However, these effects are not universal. Thus, mutating this residue at the β_1_AR does not lead to significant differences in agonist binding affinity - even in the presence of Na^+^ - suggesting the activity of this receptor is not modulated by Na^+^.^35^ However consideration should be taken, as the reported experiments were done in insect Sf9 cell membranes and using only one agonist, isoprenaline. Further functional, kinetic, and structural studies will need to be performed on additional GPCRs to determine whether insensitivity to Na^+^ ions and guanine nucleotide with highly potent agonists is a general phenomenon.

The structure of the allosteric Na^+^ site has been finely tuned to delicately balance the transition between R and R* through stabilizing GPCRs in their ligand-free state. Our results indicate that superagonists of μOR with high intrinsic efficacy may negate this physiological role of Na^+^ by binding to a Na^+^-bound receptor state, and facilitating it’s transition into an active receptor state that is able to efficiently couple to heterotrimeric G proteins. Alternatively, Na^+^ ions and guanine nucleotides by promoting the R form of the receptor speed up agonist dissociation. It is possible that very potent agonists hold the receptor in a high affinity active ternary complex due to their slow rates of dissociation from the receptor and markedly slow the ability of Na^+^ and guanine nucleotides to influence the state of the receptor. Certainly, it is known that carfentanil^36^ and BU72^37^ dissociate very slowly from the μOR, as does BWU373U86 at δOR^25^, and a slow dissociation rate may also explain why it is more difficult for naloxone to reverse respiratory depression caused by highly potent agonists like carfentanil.^38^ Furthermore, the rate of dissociation of carfentanil is much slower than that of etonitazene which could explain their different sensitivities to Na^+^ even when considering their equivalent affinities and functional activity at μOR in this study. Whatever the mechanism, the implication is that a decrease in binding affinity due to the presence of Na^+^ is not a reliable predictor of efficacy when dealing with very potent compounds in contrast with previous findings.^6,12^ Understanding the mechanisms and specific receptor-ligand interactions that mediate Na^+^ insensitivity may allow for the design of more potent GPCR agonists that are active *in vivo*, in the presence of Na^+^ and guanine nucleotide. It is important to note that this phenomenon does not occur with existing endogenous opioid peptides, but exclusively with high affinity and intrinsic efficacy ligands.

In conclusion, the influence of Na^+^ ions and guanine nucleotides on GPCR activation should not be viewed as a uniform or obligatory inhibitory mechanism. The allosteric effect of Na^+^ ions appears to be probe-dependent, varying with the chemical scaffold of the bound agonist. Different agonists, or in the case reported here, superagonists can bind to low affinity receptor and lead to the collapse of the Na^+^ binding pocket to generate active states. Alternatively, superagonists may hold the receptor in an active conformation due to their slow dissociation rates or could even stabilize unique Na⁺-bound intermediate or active-like receptor states that remain competent for G protein engagement. In this context, superagonists may exploit Na^+^ binding rather than oppose it, thereby promoting receptor activation through Na⁺-bound states that conventional agonists alone cannot access. It is possible that the role of Na^+^ in μOR activation may encompass multiple parallel trajectories, some requiring ion displacement and others proceeding through stabilized Na⁺-bound intermediates that facilitate G protein coupling.

## Abbreviations

CHO: Chinese hamster ovary;
DAMGO: [D-Ala^2^, NMe-Phe^4^, Gly-ol^5^]-enkephalin;
GDP: Guanosine 5’-diphosphate;
GPCR: G protein-coupled receptor;
GTPγ^35^S: (Guanosine-5’-O-(γ-thio) triphosphate-[^35^S]);
μOR: mu-opioid receptor.

